# The Hidradenitis Suppurativa ‘Omics Database (HS-OmicsDB)

**DOI:** 10.1101/2023.01.26.524600

**Authors:** Peter Dimitrion, Ian Loveless, Li Zhou, Qing-Sheng Mi, Indra Adrianto

## Abstract

Large scale meta-analyses of genomics and genetics have spurred research in a number of fields, such as cancer, genetics and immunology. Publicly available ‘omics databases provide valuable hypothesis generating and validation tools. To date, no such initiative has been undertaken for Hidradenitis Suppurativa (HS), an inflammatory skin disease of unknown etiology. We present here, a longitudinal initiative seeking to aggregate publicly available ‘omics data to enhance research efforts in HS. In its first iteration, we include bulk and single-cell RNA sequencing data from untreated HS patients. Our data, aggregated from publicly available GEO datasets provides a tool to profile gene expression in specific tissue types (i.e. lesional, perilesional, nonlesional and healthy skin) as well as map cell-specific gene expression on single-cell data from HS lesions.

## Introduction

Hidradenitis suppurativa (HS) is a chronic inflammatory skin disease that has among the greatest patient-reported impact on quality of life of any skin condition (Sabat et al., 2020). Patients with HS develop exceptionally painful abscesses and lesions. The prevalence of HS is estimated to be about 1%, and the condition trends with ethnic background and family history, pointing to a key role for inheritance (Sachdeva et al., 2021).

Recent studies point toward HS as an autoinflammatory keratinization disease, and a recent explosion of transcriptomic studies have identified numerous inflammatory pathways and cell-types involved (van Straalen et al., 2022). The growing list of cellular and molecular mediators in HS implies pathogenetic heterogeneity defined by a penchant for a specific immune dysregulation that may be influenced by an individual’s genetic background and environment. In fact, recent data from our lab has shown that the peripheral blood immunomes of HS patients are heterogeneous (Submitted manuscript). To date, no resources exist to reconcile this heterogeneity or elucidate the functional differences between upregulated cytokines in HS. Single-cell technologies enable resolution of cell-specific disease features and have proven fruitful in uncovering novel aspects of autoinflammatory diseases. Furthermore, meta-analyses of transcriptomic data has been proved fruitful in defining unique features of other inflammatory skin diseases (Martinez et al., 2022).

## Results & Discussion

Here we present HS ‘omics database (HS-OmicsDB, https://shiny.hfhs.org/hsomicsdb/), an initiative to integrate the growing body of ‘omics data’ collected from studies of HS. The goal is to provide a tool for the HS research community to enhance investigations into the etiology of HS specific disease features. Our database compiles publicly available bulk-RNA sequencing (Gudjonsson et al., 2020, van Straalen et al., 2022) and single cell RNA sequencing data (Gudjonsson et al., 2020, Mariottoni et al., 2021)available from studies on patients with HS deposited on the gene expression omnibus (GEO). Users can explore genes of interest to determine tissue-specific and cell-specific gene expression.

We aggregated bulk-RNAseq datasets (GSE155176, GSE151243, GSE154773) and aligned to GRCh38 using HISAT2 (Kim et al., 2019) with default parameters. Raw counts were quantified using *featurecounts* from the subread linux package (Liao et al., 2019) (Figure 1A). Currently, 71 lesional (L), 19 perilesional (PL), 29 nonlesional (NL) and 11 healthy control (HC) samples are included in this database. We incorporated two built in analyses: 1) expression of a gene of interest can be compared between L, PL, NL and HC skin; and 2) gene-correlation analysis, where all genes that significantly correlate with a gene of interest are identified by a Bonferroni-adjusted *p-value < 0*.*05* can be exported for utilization with gene enrichment tools (e.g. ingenuity pathway analysis (IPA)).

**Figure 1.**
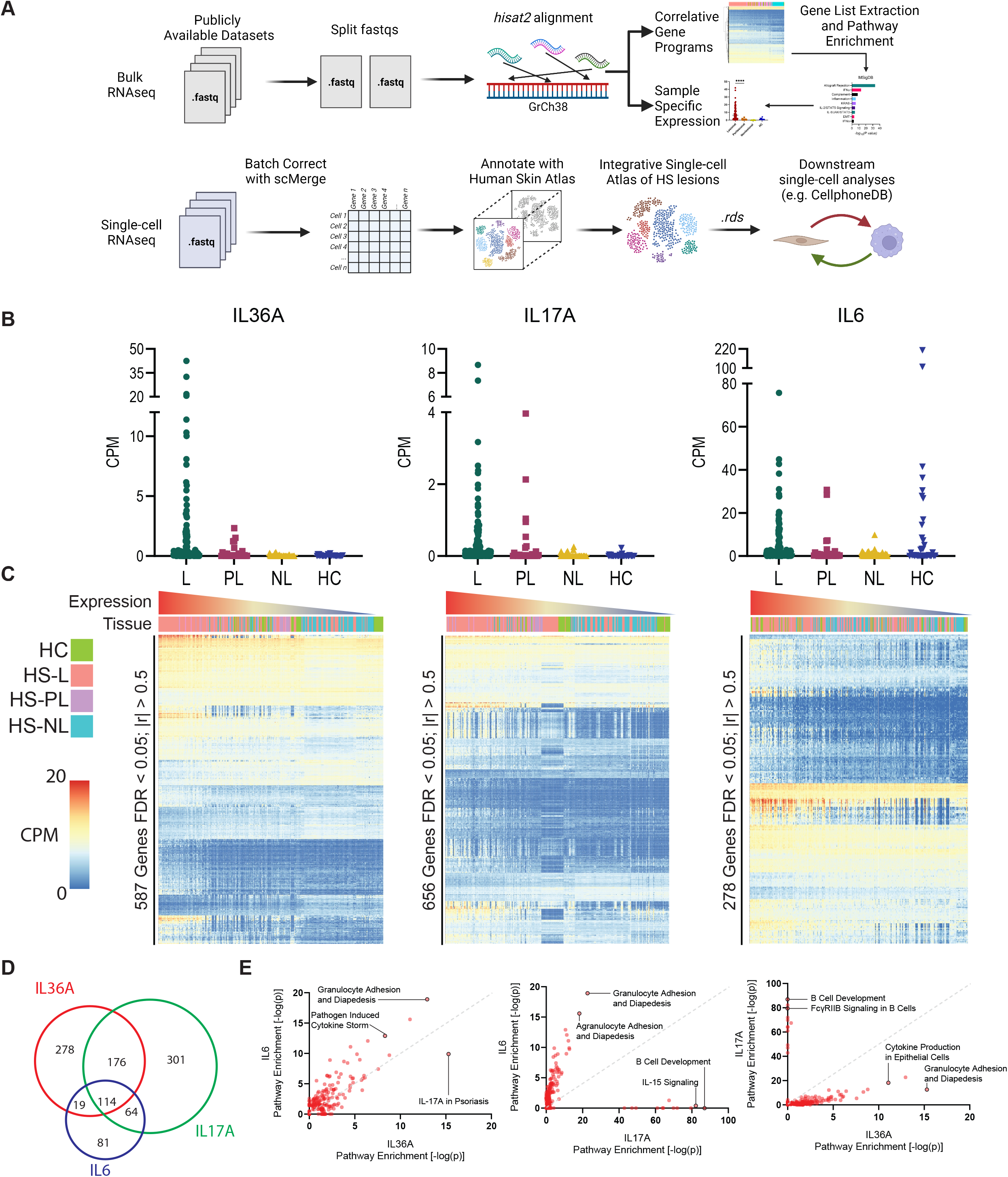
Bulk transcriptomic meta-analysis using HS-OmicsDB: (A) HSOmicsDB workflow for the aggregation of bulk and single-cell transcriptomic data. (B) Tissue-based expression of IL36a, IL17a, and IL6 in lesional (L), perilesional (PL), nonlesional (NL) and healthy control (HC) skin. (C) Heatmaps of genes that are significantly correlated with the designated cytokine. Samples are ordered based on the expression of the designated cytokine. (D) Unique and shared genes from gene-correlation analysis. (E) Pairwise comparison of pathways enriched from unique correlated genes.

Using these analyses, we compared three inflammatory cytokines: IL36, IL17, and IL6. We found all three cytokines were upregulated in HS-L over the other tissue samples (Figure 1B). Some HC skin samples had high expression of IL6. Correlation analysis found that IL36, IL17 and IL6 had 1616, 2153, and 2508 significantly correlated genes, respectively (Figure 1C). Comparing these gene lists only 570 genes were found sharing between these three gene lists (Figure 1D). IL6 and IL17 shared 483 genes, IL17 and IL36 shared 278 genes. IL6 and IL36 shared only 74 genes, highlighting divergent signaling pathways affected by these cytokines. Using IPA, we found that focal-adhesion kinase (FAK) signaling components were more enriched in IL6 and IL17 related genes. IL36 related genes were instead preferentially enriched for interferon (IFN) signaling, agranulocyte adhesion and diapedesis pathways over IL17 and IL6, respectively. These highlight the feasibility of HS-OmicsDB being used as a compelling platform for dissecting the downstream functions of specific cytokines (or any genes of interest) in HS studies. Meta-analyses such as this could support ongoing investigations unraveling the role of various effector molecules, cytokines, and chemokines in the pathogenesis of HS.

We also incorporated single-cell RNA sequencing data from two publicly available datasets (GSE154775 and GSE175990) (Gudjonsson et al., 2020, Mariottoni et al., 2021) that assayed whole skin biopsies from patients with HS. Samples aligned to GRCh38 using CellRanger v6.1.1. Finally, cells were combined, clustered, and annotated using a combination of manual annotation and label transferring from the human skin atlas using the Harmony and Seurat packages (Korsunsky et al., 2019, Reynolds et al., 2021, Stuart et al., 2019) (Figure 2A & 2B). Genes of interest are simultaneously plotted on the UMAP and violin plots alongside bulk-RNA sequencing to provide insight into expression of genes at a cell-specific level.

**Figure 2.**
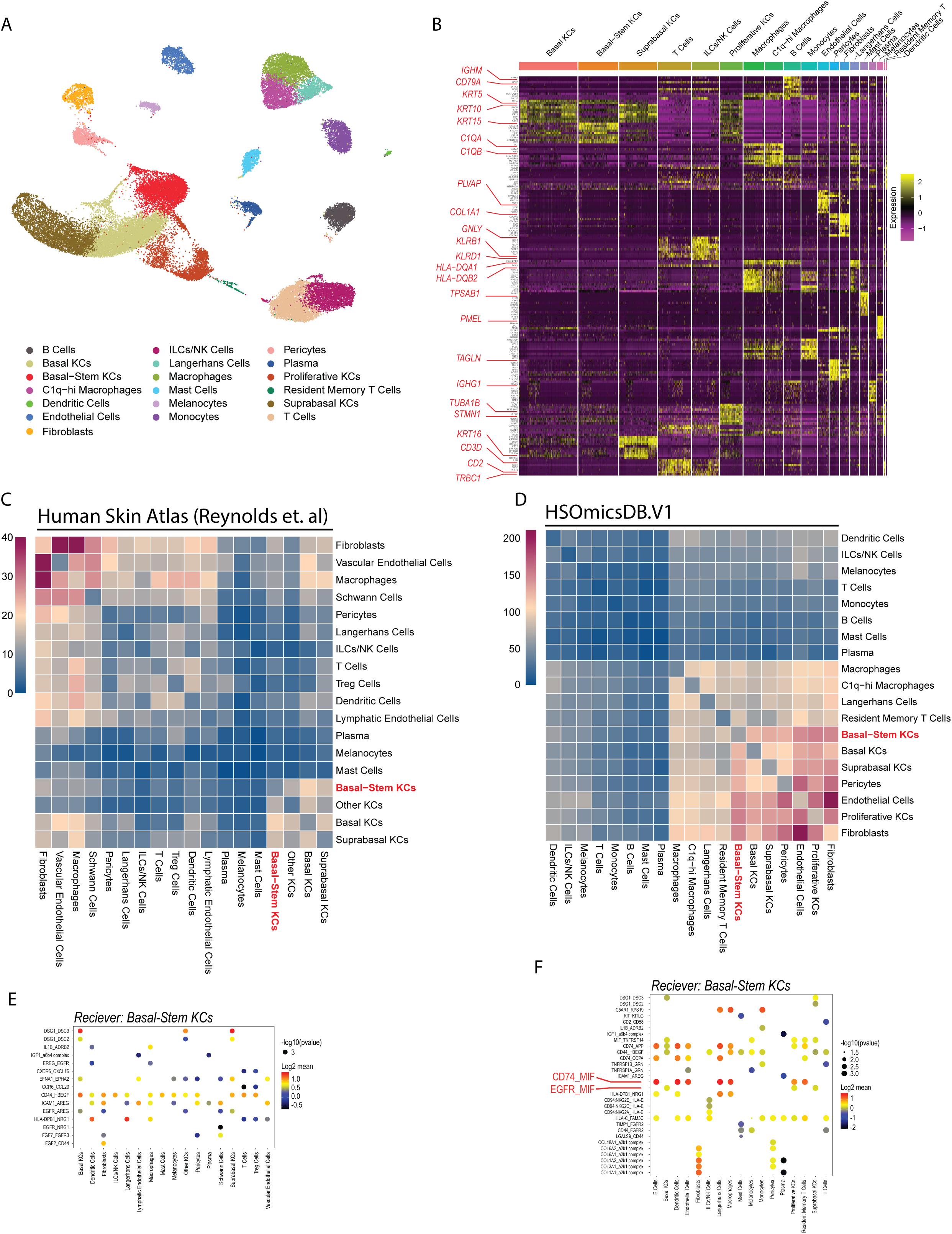
Meta-analysis of aggregated single-cell data from HS identifies dysregulated interactions poten tially involved in the formation of dermal tunnels. (A) Annotated uniform manifold approximation and projec tion (UMAP) plot of single-cell data derived from whole HS skin. (B) Heatmap of normalized gene expres sion highlighting marker genes and gene clusters for UMAP annotations. (C) Interaction frequency from CellphoneDB output from healthy control samples from the Human Skin Atlas. (D) Interaction frequency from CellphoneDB output for HSOmicsDBV.1. (E) Top five ligand-receptor interactions where Basal-stem KCs are the receiver for healthy control samples from the human skin atlas. (F) Top five ligand-receptor interactions where Basal-stem KCs are the receiver for healthy control samples from HS-OmicsDB.

Furthermore, we’ve added the ability for users to download the .*rds* file for additional single-cell applications. As an example, we employed CellphoneDB (Efremova et al., 2020) to compare ligand-receptor (L-R) interactions between HS and HC cells (from the human skin atlas) (Reynolds et al., 2021). This analysis highlighted significant dysregulation of cell-cell communications in HS skin (Figure 2C & D). Notably in HS we noticed a marked increase in the interactions between basal-stem keratinocytes (sKCs; *Krt5*^*+*^ *Krt15*^*+*^ *Krt10*^*-*^) with endothelial cells (ECs), fibroblasts, macrophages (Macs) and Langerhans cells (LCs). Further analyzing the signals received by sKCs in this dataset showed enrichment of distinct L-R pairs between HC and HS. Macrophage migration inhibitory factor (MIF) derived from ECs, Macs, and LCs are among the top 5 L-R interactions in HS and absent in HC (Figure 2E & 2F). Previous studies have found MIF is increased in psoriasiform dermatitis, where it was found to be critical for KC proliferation (Bezdek et al., 2018). In the context of HS, the increased interactions between sKCs, ECs, fibroblasts, Macs and LCs may be indicative of interactions present in dermal tunnels. Additionally, there could be a role for MIF- signaling on sKCs that may be involved in the formation of tunnels.

In the wave of thriving HS research, this initiative could serve as a valuable resource for future investigations. HS-OmicsDB will prove to be a valuable validation and hypothesis generating tool in the study of HS. Future versions will incorporate the growing number of ongoing HS studies employing ‘omics technologies (e.g. proteomics, lipidomics, mass cytometry, spatial sequencing etc.) to further enhance research in the HS community. We hope to build this resource to provide a ‘Rosetta stone’ of the complex cutaneous dysregulation that occurs in HS by integrating data from multiple ‘omics technologies, that is easy to be use and flexible in its application. Such a tool will help build an integrative view of the dynamic process that occurs throughout the progression of HS.

## Author Contributions

Conceptualization: QSM, IA, LZ; Formal analysis: IA, IL, PD; Funding acquisition: QSM, IA, LZ; Resources: IA, IL; Investigation: PD, IL, IA, QSM, LZ; Supervision: QSM, IA; Writing-Original draft preparation: PD

## Acknowledgements

This work was supported by the following grants: R01AR078688, R21AR079089, R33AR076803 to IA and Q-SM from the NIAMS/NIH; Henry Ford Immunology Program Research Support to IA, LZ, and Q-SM. We thank Gabriel Bernat for setting up and maintaining the Shiny server in our group.

## Data Availability

Data presented herein is freely accessible at https://shiny.hfhs.org/hsomicsdb/. Additional files are available upon reasonable request.

